# Horizontal gene transfer facilitates the spread of extracellular antibiotic resistance genes in soil

**DOI:** 10.1101/2021.12.05.471230

**Authors:** Heather A. Kittredge, Kevin M. Dougherty, Sarah E. Evans

## Abstract

Antibiotic resistance genes (ARGs) are ubiquitous in the environment and pose a serious risk to human and veterinary health. While many studies focus on the spread of *live* antibiotic resistant bacteria throughout the environment, it is unclear whether extracellular ARGs from *dead* cells can transfer to live bacteria to facilitate the evolution of antibiotic resistance in nature. Here, we inoculate antibiotic-free soil with extracellular ARGs (eARGs) from dead *Pseudeononas stutzeri* cells and track the evolution of antibiotic resistance via natural transformation – a mechanism of horizontal gene transfer involving the genomic integration of eARGs. We find that transformation facilitates the rapid evolution of antibiotic resistance even when eARGs occur at low concentrations (0.25 μg g^-1^ soil). However, when eARGs are abundant, transformation increases substantially. The evolution of antibiotic resistance was high under soil moistures typical in terrestrial systems (5%-30% gravimetric water content) and was only inhibited at very high soil moistures (>30%). While eARGs transformed into live cells at a low frequency, exposure to a low dose of antibiotic allowed a small number of transformants to reach high abundances in laboratory populations, suggesting even rare transformation events pose a risk to human health. Overall, this work demonstrates that dead bacteria and their eARGs are an overlooked path to antibiotic resistance, and that disinfection alone is insufficient to stop the spread of antibiotic resistance. More generally, the spread of eARGs in antibiotic-free soil suggests that transformation allows genetic variants to establish at low frequencies in the absence of antibiotic selection.

**Importance:** Over the last decade, antibiotics in the environment have gained increasing attention because they can select for drug-resistant phenotypes that would have otherwise gone extinct. To counter this effect, bacterial populations exposed to antibiotics often undergo disinfection. However, the release of extracellular antibiotic resistance genes (eARGs) into the environment following disinfection can promote the transfer of eARGs through natural transformation. This phenomenon is well-documented in wastewater and drinking water, but yet to be investigated in soil. Our results directly demonstrate that eARGs from dead bacteria are an important, but often overlooked source of antibiotic resistance in soil. We conclude that disinfection alone is insufficient to prevent the spread of ARGs. Special caution should be taken in releasing antibiotics into the environment, even if there are no *live* antibiotic resistant bacteria in the community, as transformation allows DNA to maintain its biological activity past microbial death.

## Introduction

Antibacterial resistance is a global threat to public health and could have a higher death toll than cancer by 2050 (O’Neill 2016). To reduce the impacts of antibiotic resistance on human health, we need to understand how antibiotic resistance genes (ARGs) move through the environment (Finley et al. 2013; Singer et al. 2016). However, the evolution of antibiotic resistance has traditionally been viewed as a clinical problem, and consequently little is known about when and how novel antibiotic resistant pathogens emerge from natural systems (Berglund 2015; Peterson and Kaur 2018). ARGs in the environment are particularly concerning because they pose a significant threat to food and water resources (Finley et al. 2013) and can spread to new hosts through horizontal gene transfer (HGT) (Jiang et al. 2017; Bengtsson-Palme et al. 2018; Peterson and Kaur 2018; McInnes et al. 2020). The spread of ARGs via HGT is a major mechanism in the rise of antibiotic resistance (Von Wintersdorff et al. 2016). However, the environmental variables that promote the transfer of ARGs remain poorly understood, despite well-documented instances of ARGs moving from the environment to the clinic (Poirel et al. 2002, 2005; Wright 2010; Singer et al. 2016).

An important, but often overlooked source of environmental ARGs is extracellular DNA (eDNA) (Mao et al. 2014; Dong et al. 2019). Extracellular ARGs (eARGs) enter the environment through active secretion or bacterial death, and once there can integrate into new bacterial genomes through a mechanism of HGT called natural transformation. Soil harbors one of the largest environmental reservoirs of eARGs (Heuer and Smalla 2007; Binh et al. 2008) and is home to many antibiotic producing bacteria that could select for the maintenance of ARGs in new hosts (Peterson and Kaur 2018). Since eARGs can persist in soil for more than 80 days, in comparison to less than 1 day in aquatic environments, the odds of a transfer event and subsequent spread are high in soil (Lorenz and Wackernagel 1987). Overall, understanding the occurrence of these transfer events will be important for the early detection of multi-drug resistance, especially since many antibiotic resistant pathogens are known to acquire eARGs through natural transformation (Lerminiaux and Cameron 2019).

The use of animal manures to fertilize agriculture fields can promote the spread of eARGs to the farm environment (Binh et al. 2008; Heuer et al. 2011; Ruuskanen et al. 2016). Although transformation generally increases with the availability of eDNA in laboratory studies, transformation may proceed differently in complex and spatially heterogenous environments like soil. For instance, spatial barriers have been shown to limit the transfer of plasmids via conjugation (Freese et al. 2014). Another possible barrier to transformation is the rate of eDNA decay, which can vary substantially with soil properties like water content (Sirois and Buckley 2019). Specifically, the conditions which favor eDNA stability may differ from those that favor competence – the physiological state of transforming cells. For instance, wetter soils tend to favor active growth and thus transformation but can simultaneously promote eDNA degradation (see Fig. 1 for the factors likely to affect transformation in soil). Finally, soil microbial processes like biofilm formation and cell motility are likely to affect access to eARGs and thus the efficiency of gene transfer (Molin and Tolker-Nielsen 2003; Madsen et al. 2012).

**FIG 1.**
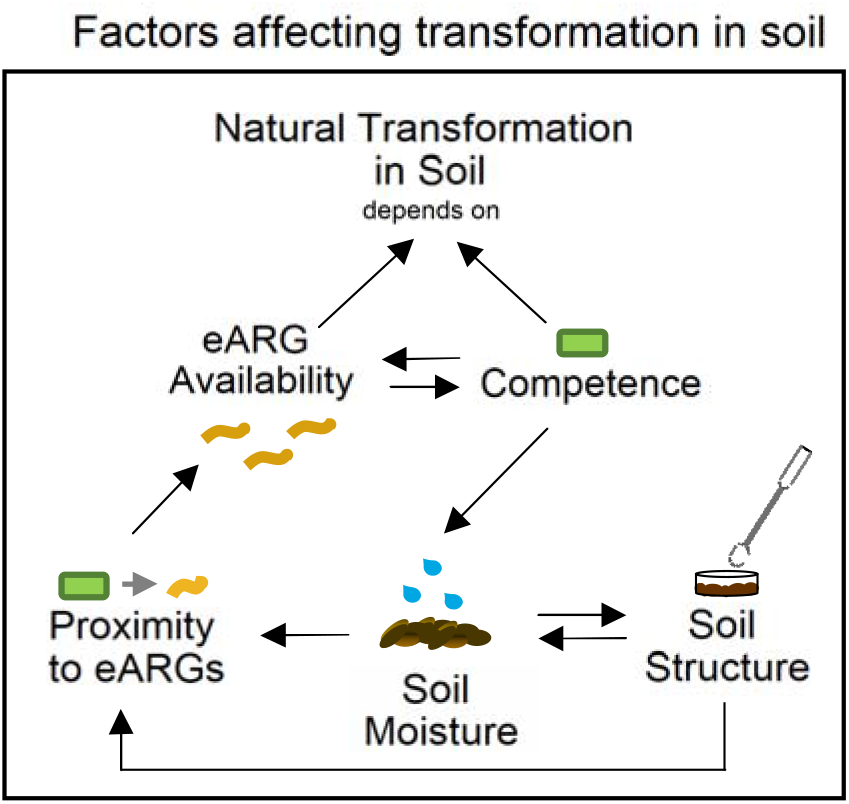
Soil characteristics likely to affect natural transformation. Transformation requires cellular competence and the presence of eARGs. But is also likely to depend on the soil moisture, soil structure, and proximity to eARGs. Arrows show possible interactions between soil characteristics and point towards the effected variable.

The presence and persistence of eARGs in agricultural soils suggests dead bacteria could be an overlooked source of antibiotic resistance in natural systems. However, the prevalence of transfer events in soil and the persistence of antibiotic resistant phenotypes has not been investigated in detail. Here, we address this knowledge gap by incoulating agricultural soils with eARGs under a series of environmentally relevant soil conditions. We then track the evolution of antibiotic resistance into soil populations of *Pseudomonas stutzeri* – a model organism for studying transformation in soil (Sikorski et al. 1998).

## Results

Inoculating eARGs into soil resulted in horizontal gene transfer through natural transformation. Live cells carrying transformed eARGs appeared in soil within 24 hours after eARG addition and evolved in the presence of just 0.25 μg eDNA g^-1^ soil, which, conservatively-estimated, is only a fraction (~1/100) of eDNA in field soil (see Pietramellara et al. 2009; Morrissey et al. 2015). Increasing the amount of eARGs increased the number of transformants in soil, thereby increasing the number of antibiotic resistant bacteria. While transformation occurred under a wide range of soil conditions, rates were highest at intermediate soil moistures (5-20%) and were strongly dependent on the stability of local eARG pools (see Fig. S1 for soil microcosm experimental design).

To test how the soil environment impacted transformation, we compared the number of transformants that evolved in soil to those that evolved on low (R2A) and high-nutrient (LB) agar petri dishes. Transformants appeared in soil and petri dishes at a similar frequency (Fig. 2A). However, it took 4 days for the frequency of transformants in soil to equal the number of transformants on agar petri dishes (Fig. 2A). While transformation initially proceeded slower in soil than under optimal laboratory conditions, the soil environment generally posed a weak barrier to transformation (24-72hrs, p<0.001 across the 3 treatments).

**FIG 2.**
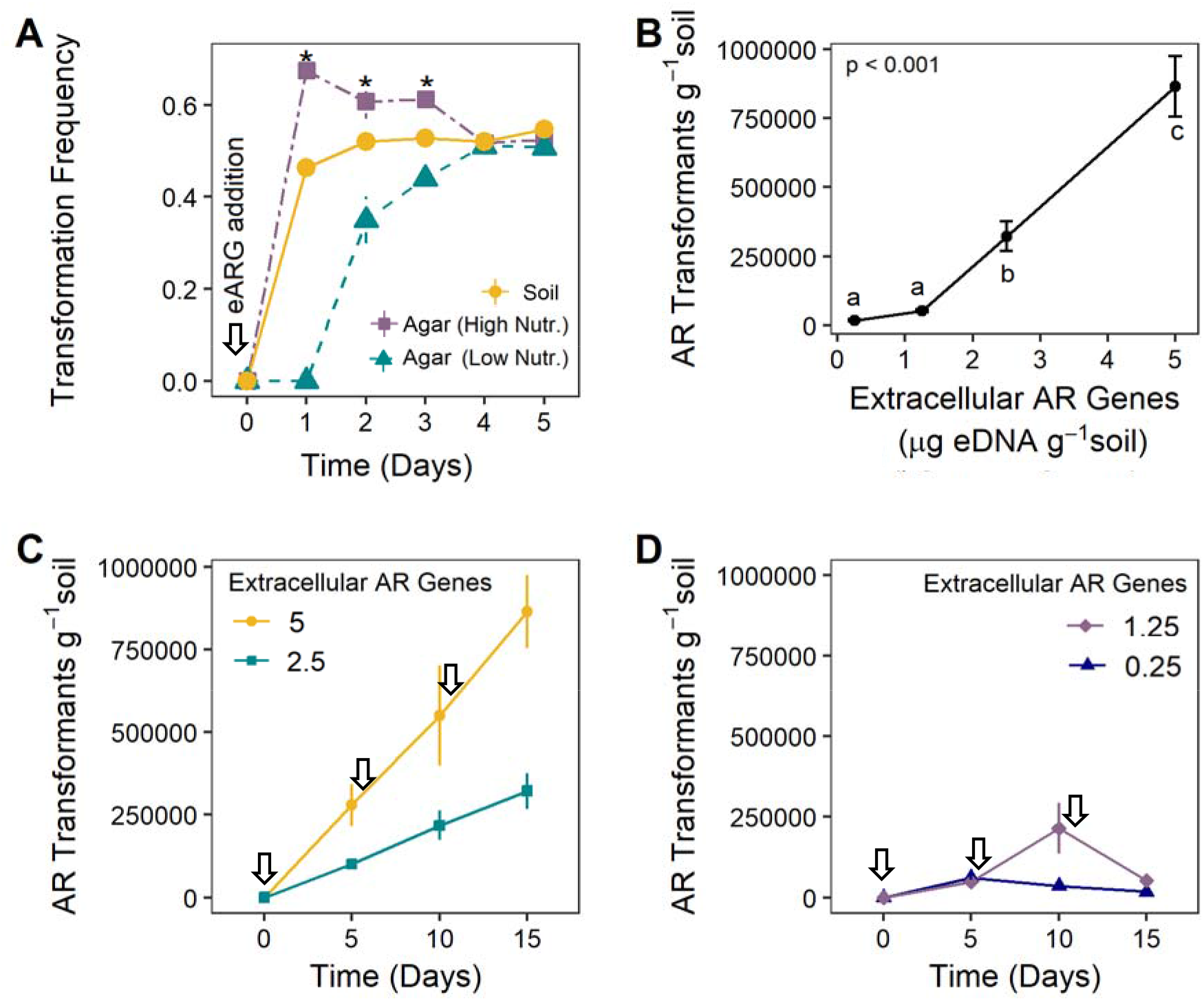
The relationship between eDNA availability and transformation. **(A)** Comparison of transformation in soil versus on petri dishes (High Nutrient = Luria Broth (LB); Low Nutrient = Reasoner’s 2A Agar (R2A)). Asterisks indicate that the transformation frequency varied across the 3 treatments on that day. **(B)** The relationship between eARG concentration and the appearance of transformants in soil (day 15 from C,D). The eDNA concentration ranged from <1% to 10% of a standard soil eDNA pool. **(C-D)** Time series tracking the appearance and maintenance of transformants when soil was supplemented with period inputs of eARGs at either, **(C)** 5, 2.5, **(D)** 0.25, or 1.25 μg eDNA g^-1^ soil. eARGs were added on day 0, 5, and 10 after counting transformants. The arrows indicate the timing of the eARG additions. (A-D) Points/bars represent the average number of transformants g^-1^ soil and error bars show the standard error of the mean (n=8 replicates).

One of the strongest controls on transformation was the availability of eARGs. We found that transformation scaled linearly with the concentration of eDNA but only in soils inoculated with at least 2.5 μg of eDNA g^-1^ soil (Fig. 2B, p<0.001). Periodic inputs of large concentrations of eDNA (>2.5 μg), increased the number of transformants by an equal magnitude (Fig. 2C). Periodic inputs of small concentrations of eDNA (<1.25 μg), did not always increase the number of transformants (Fig. 2D).

When the eDNA concentration was held constant and soil microcosms were incubated at 5, 10, 20, 30 or 40% soil moisture, we found that transformation trended highest at 10% soil moisture (though only significantly higher than at 30% and 40%, p<0.001, Fig. 3A). However, the number of transformants at 5% soil moisture did not increase until day 5 of the experiment after the second dose of eARGs (data not shown). To elucidate if biofilm formation is critical for transformation, as has been demonstrated in the lab (Fig. S2C), we homogenized soils held at 10% soil moisture. Homogenizing the soil every 2h completely prevented transformation from occurring, while homogenizing every 8h reduced the frequency of transformation events compared to a non-homogenized control (Fig 3B, p=0.03).

**FIG 3.**
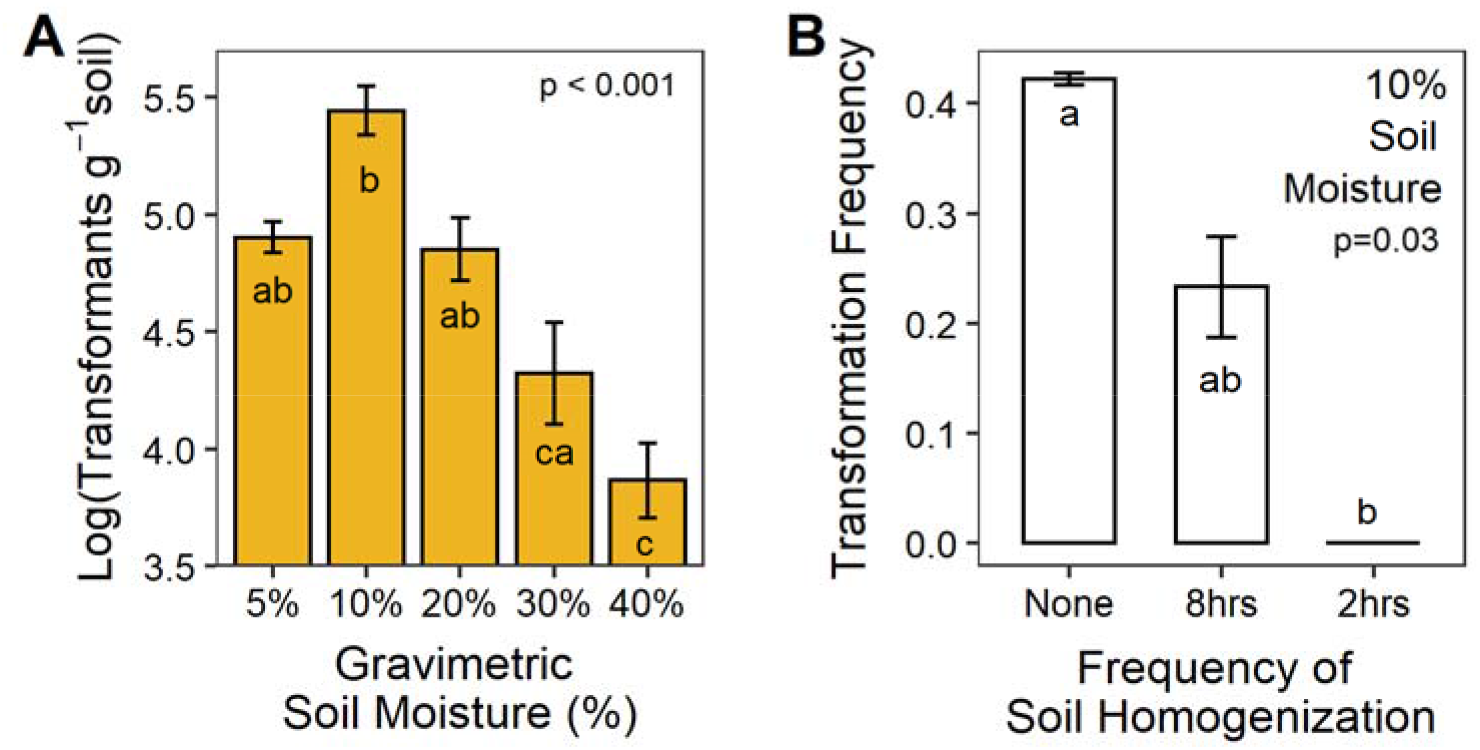
The relationship between soil moisture and transformation. **(A)** The number of transformants in soil incubated at 5%, 10%, 20%, 30% or 40% soil moisture over a 10-day experiment. **(B)** The relationship between the frequency of soil homogenization and the evolution of transformants at 10% soil moisture. Homogenization was conducted every 2hrs, 8hrs or never over a 48hr window. Bars represent the average number of log10(transformants g^-1^ soil) and error bars show the standard error of the mean (A: n=8, B: n=4 replicates).

Next, we tested the effect of separating *P. stutzeri* cells from local eARG sources and found that this spatial separation posed a substantial barrier to transformation in soils maintained at 10, 20, 30 or 40% soil moisture (Fig. 4). Even in wet soils, where dispersal to eARG sources presumably was high, there were no transformants in any of the eDNA pools (dispersal at 10% vs 40% soil moisture, p=0.0289, Fig. 4B,E). Meanwhile at 10% soil moisture there were significantly more transformants than at any other soil moisture, even though transformants only appeared in the closest eDNA pool (transformants x soil moisture, p<0.001, Figure 4B,E). Overall, dispersal and transformation happened at intermediate rates at 20% and 30% soil moisture (Fig. 4B-F).

**FIG 4.**
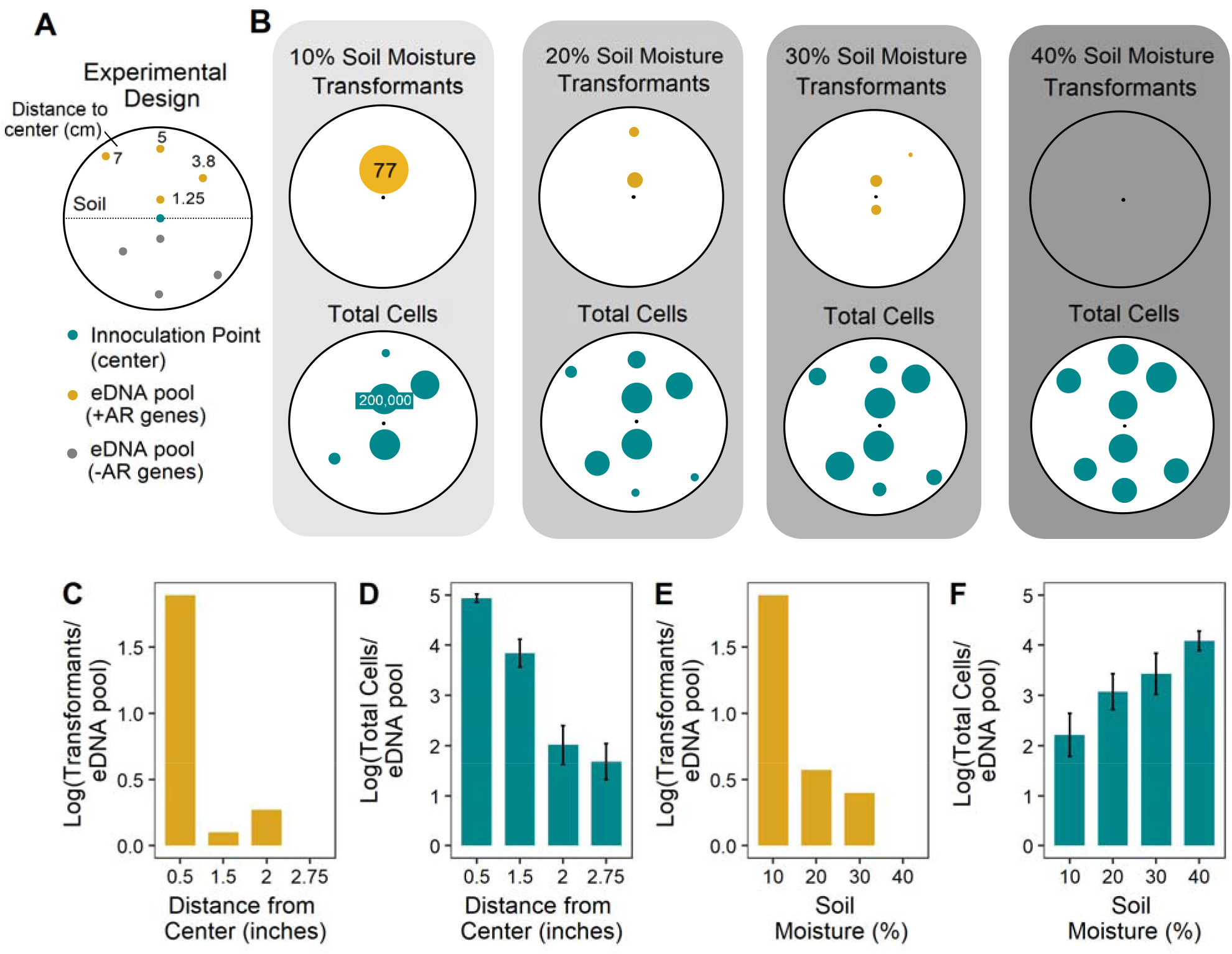
The relationship between dispersal and transformation. **(A)** Location of eDNA pools in soil microcosms set-up in 150 x 15mm petri dishes. Transformable cells were added to the center of the plate; the top 4 eDNA pools have eARGs (yellow) and the bottom 4 do not (gray). **(B)** The top panel shows the average number of transformants, and the bottom panel shows the average cell count in each eDNA pool after 5 days of dispersal. The size of the dot increases as the number of cells increases. **(C)** The total transformants and **(D)** the average cells at each distance, pooled across the four soil moistures. **(E)** The total transformants and **(F)** the average cells at each soil moisture, pooled across the four distances. (C,E) represent the sum of transformants across the replicates. Error bars show the standard error of the mean (n=4 replicates).

Lastly, to test the probability that transformed eARGs could reach high abundances in a population of *P. stutzeri*, we used a lab-based assay to compare the establishment of an equal concentration of ‘eARGs’ vs. ‘antibiotic resistant cells’ (i.e., dead vs alive cells, respectively) (Fig. 5). We compared the success of antibiotic resistant alleles in populations challenged with 10% or 25% of the lethal dose of antibiotic (gentamicin) and found that live invaders reached high frequencies in both treatments (Fig. 5, p<0.001). However, when the selective pressure increased to 25% of the lethal dose of gentamicin, both the live invader and transformed eARGs reached high abundances in the population (Fig. 5, p<0.001). Overall, antibiotic resistant transformants took 24 hours longer than live invaders to establish at high frequencies due to the time needed for transformation to transpire (Fig. 5).

**FIG 5.**
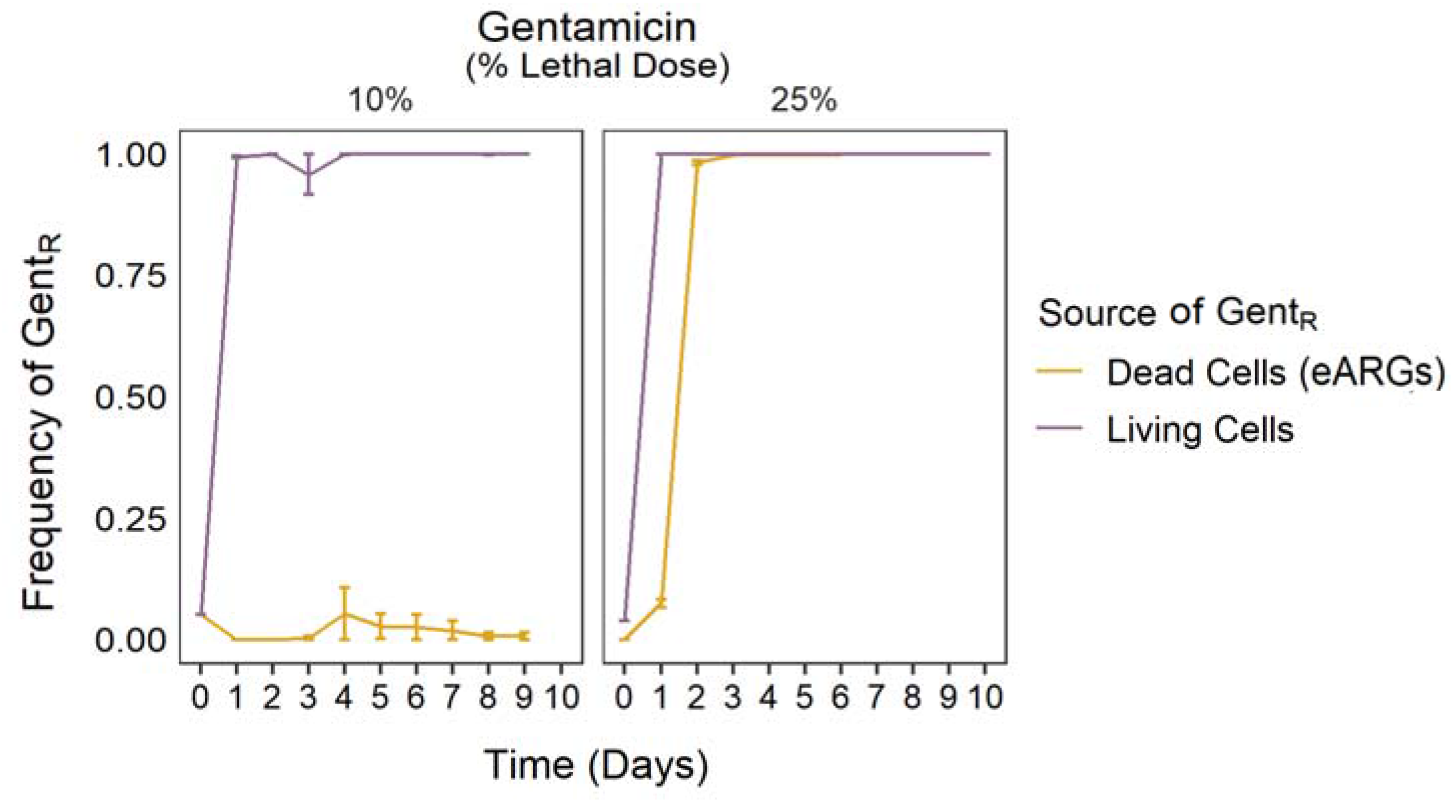
Transformation vs. invasion of antibiotic resistant genes. Frequency of gentamicin resistance (Gent_R_) over 10 days in *P. stutzeri* populations provided live or dead gentamicin resistant cells. ‘Live’ treatments started at 4% Gent_R_ (left). ‘Dead’ treatments started at 0% Gent_R_ but were provided eDNA encoding Gent_R_ (right). Error bars show the standard error of the mean (n=4 replicates).

## Discussion

In order to reduce the impacts of antibiotic resistance on human health, we need a better understanding of the ecological dimensions that promote the transfer of eARGs through natural systems (Zhang et al. 2020). In this study, we show that high concentrations of eARGs in soil increase the number of transformants, ultimately increasing the prevalence of antibiotic resistant bacteria in antibiotic-free soil. We find that transformants appear under most conditions typical for terrestrial soils (10-20% moisture), and transformation efficiency decreases at high soil moistures and with soil mixing. In addition, we find that eARGs can establish with the same success as live antibiotic resistant invaders in laboratory populations challenged with a low dose of an antibiotic (25% the lethal dose). Overall, the sustained biological activity of eARGs, even after bacteria death, suggests eARG removal should be incorporated into plans to combat antibiotic resistance.

Several studies have now posited that the spread of antibiotic genes into diverse bacterial lineages occurs via widespread HGT (Martinez 2008; McInnes et al. 2020). Here, we show that eARGs supplied by dead bacteria are readily transformed into soil bacteria, with the potential for HGT scaling with the abundance of eARGs (Fig. 2B,C). Our findings provide novel evidence that large concentrations of environmental eARGs can drive the evolution of antibiotic resistance. Thus, this information should be incorporated into our approach to combating antibiotic resistance in the environment. For instance, many disinfection methods focus on killing live bacteria, but may be more effective if they also reduce eARG pools, which we find to be equally effective at disseminating antibiotic resistance. This may explain why practices like composting manure prior to application on agricultural fields has been found to both increase and decrease the occurrence of ARGs depending on the native bacterial community (Ezzariai et al. 2018) and other soil conditions (Qian et al. 2018). Interestingly, manures composted at high temperatures, which promotes the degradation of eDNA, can be most effective in reducing ARGs (Li et al. 2017), supporting our findings that DNA degradation is a critical factor in reducing environmental concentrations of eARGs.

Despite the dangers of low-levels of eARGs persisting in soil for an extended time, the fate of most extracellular DNA is likely degradation (Lorenz and Wackernagel 1987). While low levels of eDNA can persist in soil for 80+ days, previous work indicates that 99% of eDNA is degraded in the first ~7 days in soil (Sirois and Buckley 2019; McKinney and Dungan 2020). Consequently, the most important role of soil conditions in regulating transformation may be the effect of moisture on the rate of eDNA decay, and likely explains our finding that transformation declined at higher soil moistures (Fig. 3A) even though recipient cells had access to more eARGs (Fig. 4B). Similar trends have been observed for the soil bacterium *Acinetobacter calcoaceticum* whose transformation frequency was highest at 18% soil moisture, but lowest at 35% soil moisture (Nielsen et al. 1997). However, direct comparisons between *A. calcoaceticum* and *P. stutzeri* are difficult because *A. calcoaceticum* transformation efficiency was only measured above 18% soil moisture. Consequently, our work provides surprising evidence that transformation is favored at lower soil moistures, countering the idea that many microbial processes increase with soil moisture (Trivedi et al. 2016).

The relationship between soil moisture and transformation could vary widely across bacterial species though and is likely to depend on how individual taxa regulate transformation. Additionally, the agricultural soils used in this study rarely exceed 30% soil moisture (Fig. S3), when transformation was lowest in our study (Fig. 3A). A biological explanation of this is that soils are optimized to support HGT in their native range of soil moistures. Alternatively, the frequency of eARG transformation may be explained by soil physical factors. An interesting follow-up study would be to quantify transformation in wetland or sediment soils, where water content is generally higher.

We also observed that soils held at 10% soil moisture only supported high levels of transformation if they were not disturbed by physical homogenization, suggesting an interaction between soil moisture and the physical structure of soil (Fig. 3B). Because HGT is high in biofilm communities but rare in planktonic communities (Fig. S2C), physical homogenization of the soil likely prevented the formation of mature biofilms, and subsequently prevented the spread of eARGs (Ding et al. 2019). In fact, biofilms are generally more common in dry soils where they increase microbial survivorship (Lennon and Lehmkuhl 2016), further supporting the possibility that biofilm formation is key for the horizontal dissemination of eARGs, particularly in drought-exposed soils (<10% soil moisture). Future studies could use fluorescent proteins or confocal laser scanning microscopy to better quantify the relationship between biofilm establishment and transformation efficiency in soil (Wilson et al. 2017).

A major concern in the fight against antibiotic resistance is the presence of antibiotics in the environment, as they provide a selective pressure for transformed eARGs to reach high abundances in a community. Our results highlight this danger, as laboratory populations demonstrated high levels of antibiotic resistance, whether the antibiotic resistant alleles were introduced as live or dead cells. Most alarming though, was that the success of transformants depended on the concentration of antibiotics in the surrounding environment (Fig. 5). An important future research direction will be determining the antibiotic concentrations at which eARGs can establish in a population of soil bacteria. To answer this question will require an increased understanding of antibiotic concentrations in soil and their effect on bacterial communities.

Taken together, our study reveals the most important variables for understanding the transmission of eARGs in soil and sets the stage for future experiments to scale up estimates of transformation to the whole community. Here, we used a sterile soil system, inoculated with a single bacterium to prevent competitive interactions, and to ensure the soil was antibiotic-free. However, transformation is likely lower in multi-species communities, where competitive interactions could limit access to eARGs and limit the success of transformants. Certain soil types may further alter transformation rates, and future studies could probe this relationship. Regardless, this work provides novel evidence that eARGs from dead bacteria are an overlooked, but important route in the emergence of antibiotic resistance. Specifically, we find that the availability of eARGs drives the evolution of antibiotic resistance, and that transformation is prevalent under a wide range of soil conditions – only decreasing at very high soil moistures and in homogenized soils. We recommend special caution if releasing antibiotics or eARGs into soil, as transformation allows antibiotic resistant genes to maintain their biological activity past death. Together, this work provides novel *in situ* evidence that HGT is an evolutionary force that facilitates the spread of non-selected antibiotic resistance genes in soil, which can later be selected for in the presence of antibiotics.

## Methods

### Site and soil collection

Soil cores (10cm depth by 5cm diameter) were collected in October 2018 and April 2019 from the Great Lakes Bioenergy Research Center (GLBRC) scale-up fields located at Lux Arbor Reserve Farm in southwest Michigan (42°24’ N, 85°24’ W). Plots were established as perennial switchgrass monocultures (*Panicum virgatum L*) in 2013, and before that were in a corn– soybean rotation for more than 10 years. The soils developed on glacial outwash and are classified as well-drained Typic Hapludalf, fine-loamy, mixed, mesic (Kalamazoo series) or coarse-loamy, mixed, mesic (Oshtemo series) or loamy sand, mixed, mesic (Boyer series) (Bhardwaj et al. 2011). Soils were sieved at 2mm and autoclaved in two cycles (60 minutes at 121°C; gravity cycle) separated by a 24-hr window to target dormant and spore-forming cells resuscitated during the first autoclave cycle.

### Bacterial cultures and extracellular DNA (eDNA)

Soil microcosms were inoculated with *Pseudomonas stutzeri*, strain 28a24 (Smith et al. 2014). Prior to inoculation, the bacterial cultures were grown at 30°C on an orbital shaker at 120 rpm for 24hrs in liquid luria broth (LB) media to a concentration of 10^6^ CFU/mL. All LB media used throughout the experiment followed a recipe of 10% tryptone, 5% yeast extract, and 5% NaCl (solid media contained 1.5% agarose). Stocks of antibiotic resistant extracellular DNA (eARGs) were made from a mutant *P. stutzeri*, strain encoding a gentamicin resistance gene and a LacZ gene (Tn7 transposition of pUC18-mini-Tn7T-Gm-lacZ into strain 28a24, see Choi and Schweizer 2006). eDNA was also made from the wildtype *P. stutzeri* to act as a negative control. The gentamicin resistant *P. stutzeri* cells were genetically identical to the wildtype *P. stutzeri* cells, except for the presence of the antibiotic resistance gene. The batch cultures for eDNA stocks were prepared under the same conditions specified above but were grown for 48hrs and then resuspended in sterile nanopure water. The cells for eDNA stocks were killed via heat shock (90°C for 1hr) and confirmed dead by plating. The final concentrations of eDNA ranged from 25-50 ng and were appropriately diluted for each experiment (determined using Qubit fluorometric quantification and Invitrogen Quant-iT PicoGreen dsDNA Assay Kit). *P. stutzeri’s* transformation efficiency plateaus at ~5ng eDNA under standard laboratory conditions (Fig. S2A).

### Soil microcosms

Soil microcosms were established in small 60 x 15mm petri dishes using 10 grams of dry, sterile, switchgrass soil. Except for the experiment in Figure 4, which used large 150 x 15mm petri dishes filled with 100 grams of soil. On day 0 of each experiment, the center of the microcosm was inoculated with 2mL of wildtype *P. stutzeri* cells suspended in liquid LB at a concentration of 10^6^ CFU/mL. Immediately after adding cells, we inoculated the soil with eDNA encoding gentamicin resistance genes. To control for contamination or evolution of gentamycin resistance via mutation, two additional treatments were included in every experiment; 1) 5μg of eDNA g^-1^ soil made from gentamicin susceptible *P. stutzeri* cells, and 2) sterile water without eDNA. Since transformants never appeared in the control treatments the results are not shown.

All microcosms were maintained at ~23°C and the soil was never mixed unless directly specified (e.g. in Fig. 3B). All microcosms were initially inoculated to ~40% soil moisture on day 0, and then dried to 20% soil moisture (except in the experiments manipulating soil moisture where the soil was dried according to the treatment-level soil moisture). In each experiment, we counted the number of transformants, and the population size every 5 days, and then added more eDNA to simulate periodic inputs of eARGs. After eDNA additions, soils were gradually dried back to 20% soil moisture.

### Manipulation of eARG concentration in soil microcosms

To establish a baseline for transformation in the soil system, we quantified transformation every 24hrs for 5 days after inoculation with 5μg eDNA g^-1^ soil. We ran parallel assays on petri dishes using LB and R2A (DF1826 Fisher Scientific) using the same concentration of eARGs, with each treatment consisting of 8 replicates (Fig. 2A).

To understand the relationship between the availability of eDNA and transformation, we varied the concentration of eDNA in soil between 5, 2.5, 1.25, 0.25 μg of eDNA gram^-1^ soil (Fig. 2B,C,D). We used these concentrations as they conservatively represent ~10%, 5%, 2.5% and 0.5% of the total eDNA pool in soil (Morrissey et al. 2015). The concentration of eDNA never exceeded 10%, to account for the fact that only a small percentage of soil eDNA is likely to encode antibiotic resistance genes. We ran the experiment for 15 days and added eARGs to soil on day 0, 5, and 10 of the experiment, with each treatment consisting of 8 replicate soil microcosms.

### Manipulation of soil moisture in soil microcosms

To determine how soil moisture affected transformation, we maintained soil microcosms at 5, 10, 20, 30 or 40% gravimetric soil moisture over a period of 10 days (Fig. 3A, i.e. soil moisture = [weight after water addition – dry weight] / dry weight]*100). In this experiment, all the microcosms were inoculated with an intermediate concentration of eDNA (2.5μg g^-1^ soil) and eARGs were added on day 0 and 5. We report the number of transformants present on day 10, using 8 replicate microcosms per treatment. To understand if the physical structure of the soil was important for transformation, we manipulated the physical structure of the soil by mixing the soil every 2hrs, 8hrs or never, throughout a 48hr period (Fig. 3B). Each treatment consisted of 4 replicates and the eDNA concentration was maintained at 5 μg g^-1^ soil, with the soil moisture remaining constant at 10%. Individual microcosms were gently mixed for approximately 30 seconds using a sterile spatula at the designated intervals.

To determine if dispersal to eARGs and subsequent transformation events vary under different soil moistures, we established 8 pools of eDNA in large microcosms maintained at 10, 20, 30 or 40% soil moisture (Fig. 4A for design). Half of the eDNA pools contained eARGs (dead gentamicin resistance *P. stutzeri*) and the other half did not (dead wildtype *P. stutzeri*), acting as a control for the movement of eARGs throughout the microcosm. The 8 pools were located 1.25, 3.80, 5 or 7 cm from the center of the plate, and each pool was inoculated with 2μg eDNA g^-1^ soil. Meanwhile, *P. stutzeri* was only added to the center of the microcosm. Following inoculation, we confirmed that there were no transformants outside of established eDNA/eARG pools. The experiment ran for 5 days, using 4 replicate microcosms per soil moisture. At the end of the experiment, soil was collected from the center of each eDNA pool to count the number of transformants and total cells.

### Antibiotic resistance genes in live vs. dead cells

In a final laboratory experiment, we tested how an equal concentration of ‘eARGs’ and ‘live antibiotic resistant cells’ establish in populations of *P. stutzeri* challenged with a low dose of antibiotic (Fig. 5). Initially, we established two equal populations of *P.stutzeri* cells (DAB837 in Romanchuk et al. 2014). To one of the two populations, we added 60,000 ‘live’ gentamicin resistant *P. stutzeri* cells (Gent_R_). To the other population, we added 60,000 ‘dead Gent_R_’ cells which provided a source of eARGs. Therefore, on day 0 of the experiment, the two treatments contained 4% Gent_R_ cells and 0% Gent_R_ cells, respectively. Populations were founded in 1mL of LB media, and the experiment ran for 10 days, with each population receiving 1 mL of fresh LB media every 24hrs. Each day we counted the total number of cells on solid LB media (no antibiotic) and the number of Gent_R_ cells on solid LB media with gentamycin (50 μg/ml) + Xgal (20 μg/ml). We report the frequency of Gent_R_ genotypes (Gent_R_ cells/total cells).

### Counting Transformants

To determine the number of transformants in each soil microcosm, we weighed out 0.2g of soil from each microcosm or eDNA pool and placed it into a 1.5mL centrifuge tube. To each tube, we added 180μl of liquid LB and vortexed for 10 seconds (~10^-1^ dilution). After allowing the soil to settle for 10 minutes, we transferred the supernatant to a 96-well plate and diluted out to 10^-6^ or 10^-9^ depending on the experiment and the expected number of cells. In the experiments that manipulated eDNA concentration and soil moisture, we plated 50μl cell suspensions. For the remaining experiments we plated 10μl dots. All plating was done on petri dishes with solid LB (to count the total population size) or solid LB + gentamycin (50 μg/ml) + Xgal (20 μg/ml) to count transformants in soil. Plates were incubated at 30°C and the number of colonies counted after 48-1 72hrs. The number of cells is reported g^-1^ soil, except in Fig. 4 where it is reported per eARG pool (0.2g soil) and calculated according to the following equation: Cells per unit = Transformants μl^-1^ x [Soil slurry volume (200μl) / Soil Mass in slurry (g)].

### Statistical analyses

Prior to analyses all data were verified to meet assumptions of normality and homogeneity of variance. Data that did not conform to assumptions of homogeneity of variance were log transformed when appropriate. The results from the soil microcosm studies were analyzed by either one-way or twoway ANOVA followed by Tukey’s post hoc with test variable (i.e. soil manipulation and sampling day) as a fixed effect using the R *stats* package (R core team 2018). Experiments with multiple sampling days were analyzed by two-way ANOVA, except in certian instances, when the test variables were analyzed individually by sampling day (e.g.Fig. 3A). Results from the soil microcosm experiment in (Fig 4) were analyzed by two-way ANOVA with the ‘distance to eARGs’ and ‘soil moisture’ as fixed effects. When signficant, interactions between test variables were included in the model. The frequency of antibiotic resistant genotypes present in each laboratory population at the end of the experiment were compared using two-way ANOVA with the treatment (Live vs Dead cells) and selection regime (10 or 25% lethal dose gentamicin) as fixed effects (Fig. 5). Differences between all test variable groups were considered significant at α ≤ 0.05.

## Acknowledgements

Support for this research was provided by the DOE Great Lakes Bioenergy Research Center (DOE BER Office of Science DE-SC0018409 and DE-FC02-07ER64494), and by the NSF Long-Term Ecological Research Program (DEB 1832042 and 1637653) at the Kellogg Biological Station. This work took place on occupied Anishinaabe land where Hickory Corners, Michigan, is now located. We thank local communities and the state of Michigan for maintaining and allowing access to the field sites. This is Kellogg Biological Station contribution number 2299. Additionally, we thank David Baltrus for constructing and sharing the *Pseudomonas stuzeri* strains used in this study.

## Author Contributions

HK, KD and SE conceptualized the experimental design for the soil microcosms. HK performed all soil microcosm incubations and laboratory work. HK and SE performed the analyses and wrote the manuscript. HK, KD, SE provided comments on the manuscript.

---

**FIG S1.**
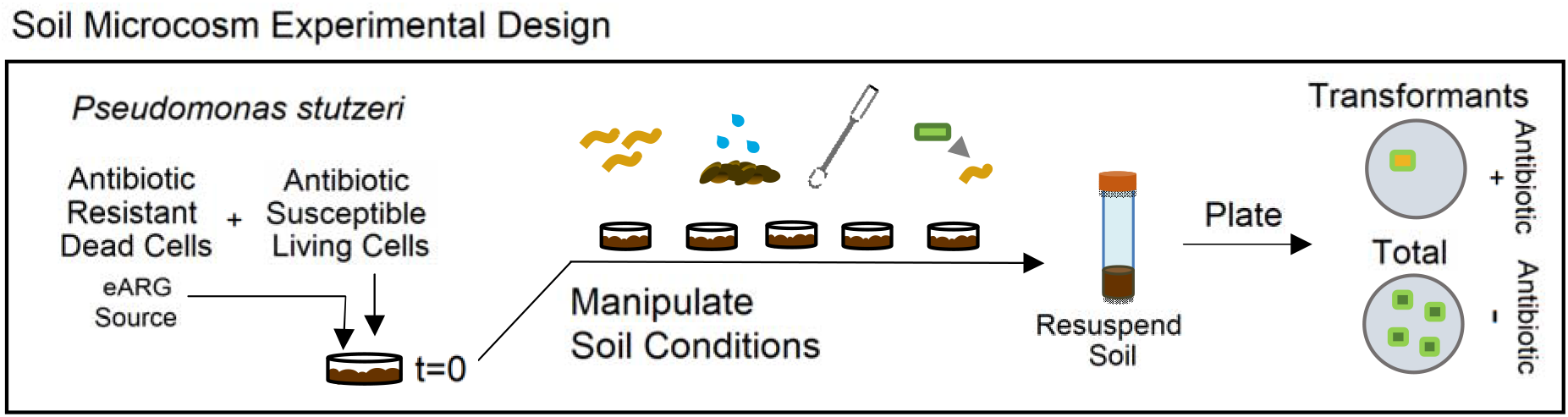
General experimental design for soil microcosms. At time zero soil was inoculated with live antibiotic susceptible *P. stutzeri* and dead antibiotic resistant *P. stutzeri* to provide a source of eARGs. Soils were then exposed to a variety of soil conditions and the number of transformants periodically counted by resuspending soil in a slurry and plating onto selective media.

**FIG S2.**
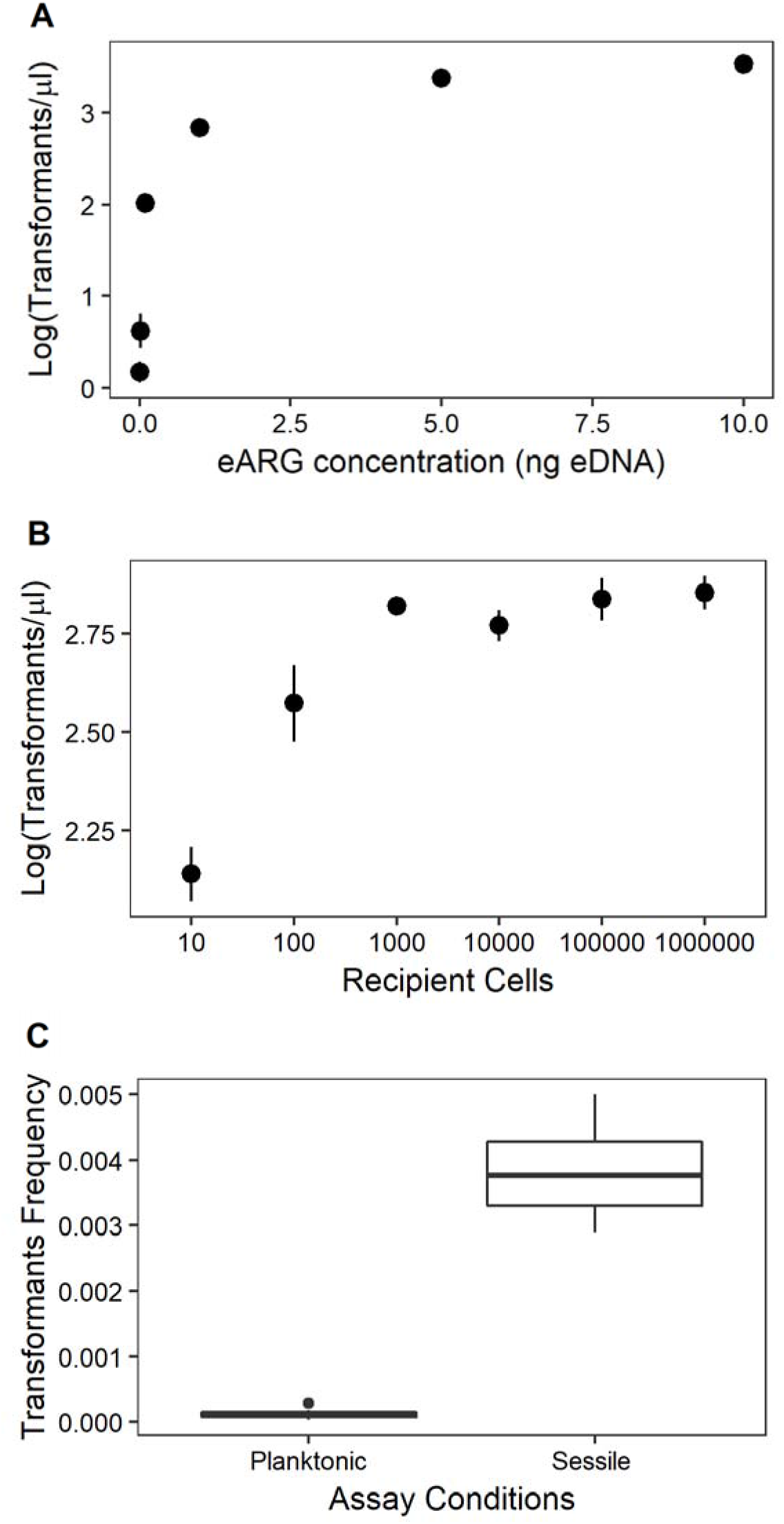
Transformation assays under laboratory conditions. **(A)** The relationship between transformation and the concentration of eDNA under laboratory conditions (0.001, 0.01, 0.1, 1, 5, 10 ng/μl eDNA). **(B)** The effect of total population size on the number of transformants. (C) Comparison of transformation in sessile (biofilm or surface attached) communities vs. planktonic communities.

**FIG S3.**
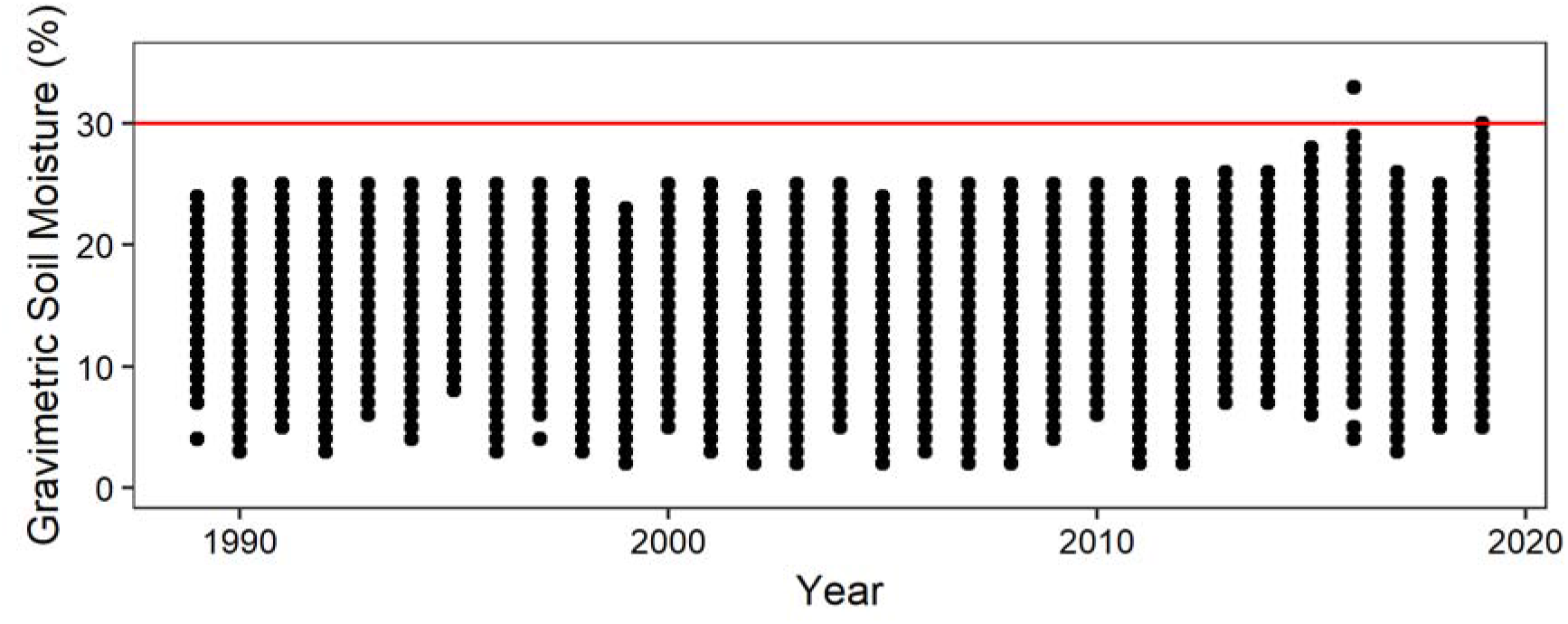
Gravimetric soil moisture from 1989 to 2019 Southwest, MI (regular measurements from April-November). Data from the Kellogg Biological Station Longterm ecological research center main cropping system. The red line shows 30% soil moisture and represents the soil moisture where transformation starts to decline in our experiment.

